# Big data from small tissues: extraction of high-quality RNA for different plant tissue types during oilseed *Brassica* spp. seed development for RNA-Sequencing

**DOI:** 10.1101/2019.12.20.885012

**Authors:** Laura Siles, Peter Eastmond, Smita Kurup

**Affiliations:** Department of Plant Science, Rothamsted Research, Harpenden, Hertfordshire, AL5 2JQ, UK

## Abstract

Obtaining high-quality RNA for gene expression analyses from different seed tissues is challenging due to the presence of various contaminants, such as polyphenols, polysaccharides and lipids which interfere with RNA extraction methods. At present, the available protocols for extracting RNA from seeds require high amounts of tissue and are mainly focused on extracting RNA from whole seeds. However, extracting RNA at the tissue level enables more detailed studies regarding tissue specific transcriptome during development. Seeds from heart stage embryo to mature developmental stages of *Brassica napus* and *B. oleracea* were sampled for isolation of the embryo, endosperm and seed coat tissues. Ovules and gynoecia wall tissue were also collected at the pre-fertilization stage. After testing several RNA extraction methods, E.Z.N.A. Plant RNA Kit and Picopure RNA Isolation kit extraction methods with some modifications, as well as the use of PVPP for seed coats and endosperms at green stages, resulted in high RNA concentrations with clear 28S and 18S bands and high RIN values. Here, we present efficient and reliable RNA extraction methods for different genotypes of *Brassica* spp for different tissue types during seed development. The high-quality RNA obtained by using these methodologies is suitable for RNA-Sequencing and gene expression analyses.

## Introduction

Oilseed rape is a crucial source of oil and protein world-wide, being a crop with high economic importance. *Brassica* spp seeds are rich in polyphenols, polysaccharides, proteins and lipids which interfere with or degrade the RNA during RNA extraction procedures. Polyphenols limit RNA extraction as they bind to nucleic acids (Mattheus et al., 2003), and polysaccharides co-precipitate with RNA (Singh et al., 2003). Removing these contaminants is a critical step, as RNA with high purity and integrity is critical for gene expression analyses. Although there are several methods available for seed RNA extraction (Kanai et al., 2017; Pereira et al., 2017; Mornkham et al., 2013; Suzuki et al., 2004); fewer protocols are available for the extraction of good quality RNA from specific seed tissue types, especially for the seed coat (Jiang et al., 2019; Liao et al., 2019; Hong et al., 2017; Kour et al., 2014). Moreover, the main constraint of these protocols is the large amount of sample required, ranging between 50 and 200 mg of tissue, in order to obtain the quality and concentration of RNA required for downstream analyses.

Since the availability of Next Generation Sequencing (NGS) methods, we have the potential to measure the level of transcripts with high quality and accuracy. With the reduction in costs for NGS methods, we can now obtain large amounts of sequence data for a large number of samples in a very cost-effective manner, providing more comprehensive studies across tissues and developmental stages.

Embryos from the Brassicaceae family follow a well-studied developmental timeline starting from globular, heart, torpedo, green and mature embryo and seed stages (Harada 1999). Herein we describe different efficient methods for isolating high-quality RNA from different tissue types from *Brassica napus* and *B. oleracea* during seed development for subsequent RNA-Sequencing analysis by only using 2-3 mg of seed coat and green stage endosperm (GSEnd) ground tissue and a small number of embryos. The implementation of these methodologies not only reduces the hours of labour intensive dissections, but also enables more detailed studies characterizing different seed tissues and the prediction of gene regulatory networks across seed development.

## Material and methods

### Plant growth conditions and tissue type collection

*Brassica* plants were grown under controlled environment conditions with a 16h photoperiod at 18°C and 15°C day and night temperatures respectively with 70% of humidity day and night. Perforated bread bags (150 mm x 700 mm, WR Wright & Sons Ltd, Liverpool, UK) were used to enclose inflorescences to prevent cross-pollination from neighbouring plants. Flowers were manually pollinated, and seeds were collected at heart, torpedo, green and mature embryo stages. Although flowers were tagged to determine the developmental stage of the seed, the embryos within the seed were visually checked to ensure they were at the correct stage of development prior to sampling. Buds were also collected 24 hours before anthesis to obtain ovules and gynoecia wall tissue pre-fertilization (Figure 1). All sampling was carried out on a clean, sterile glass slide placed on a bed of ice (dry ice for heart stage) in a 9 ml petri dish to ensure samples were cold throughout the extraction procedure. Embryos, seed coats and endosperms were separated from each other for every developmental stage except for the mature stage, in which the endosperm and the seed coat were maintained together. Gentle manipulation with forceps allowed the dissection of these tissues using a dissecting microscope. For the heart and torpedo stages the embryos were aspirated with the help of 1.5 µl RB buffer from E.Z.N.A. Plant RNA kit (Omega Bio-tek Inc., Norcross, Georgia, USA) using a P10 pipette tip for transfer to 1.5 ml microfuge tube. During *Brassica* spp seed development, the endosperm goes through a transition from syncytial to cellular phase. When the embryo is at the heart stage, the endosperm is in the liquid syncytial phase, at the torpedo stage the endosperm starts to cellularize, finally forming a single layer of cellularised endosperm by the green embryo stage (Brown et al., 1999). For heart stage endosperm (HSEnd), the seed was held gently with a pair of forceps. Next the seed coat was perforated carefully with the aid of forceps or syringe and the endosperm was collected using a P10 pipette while gently squeezing the embryo (see Supplementary Video 1). At the torpedo stage the seed was cut open, and 1 µl of RB buffer was added to facilitate removal of the endosperm with a pair of forceps. At the green stage the drop of buffer was omitted, and the single layer of endosperm was transferred using a pair of forceps and transferred to a 1.5 ml microfuge tube. Samples were dissected and immediately freeze frozen in liquid nitrogen and stored at −80°C for total RNA isolation. However, a slight modification was applied for ovules, heart and torpedo stage embryos (HSE and TSE, respectively). These samples were dissected, collected in RB buffer and ground with the help of a micro-pestle before freezing in liquid nitrogen. This methodology allowed the collection and grinding of small tissues without RNA degradation. Gynoecia walls, seed coats from all developmental stages, GSEnd, and green and mature stage embryos (GSE and MSE, respectively) were finely powdered by grinding in liquid nitrogen in a mortar and pestle. For HSEnd and torpedo stage endosperm (TSEnd), no grinding was performed as the endosperm is primarily liquid at both these stages of seed development. Based on the tissue type of interest and the developmental stage of the seed, varying amounts of starting seed material may be required. For smaller embryos as in the heart stage, approximately 100 seeds were required for efficient and high-quality RNA extractions. On the other hand, when the embryos are larger, as in torpedo, green and mature stages, only 50 seeds were needed. See Table 1 as a guideline for tissue amounts required at different developmental stages.

**Table 1:** Amount of ground tissue used for RNA extractions for different *Brassica napus* and *B. oleracea* tissue types. * = 100 HSE were ground in 250 µl of RB buffer from E.Z.N.A. Plant RNA kit and an aliquot of 20-25 µl was used for RNA extraction. An aliquot of 10-15 µl was used for RNA extraction of 50 TSE.

| Tissue type | Amount of tissue |
| --- | --- |
| Gynoecia wall | 20 mg |
| Ovules | 6 mg |
| Heart stage embryo (HSE) | 20-25 μl* |
| Heart stage endosperm (HSEnd) | 100 endosperms |
| Heart stage seed coat (HSSC) | 2-3 mg |
| Torpedo stage embryo (TSE) | 10-15 μl* |
| Torpedo stage endosperm (TSEnd) | 50 endosperms |
| Torpedo stage seed coat (TSSC) | 2-3 mg |
| Green stage embryo (GSE) | 50 mg |
| Green stage endosperm (GSEnd) | 2-3 mg |
| Green stage seed coat (GSSC) | 2-3 mg |
| Mature stage embryo (MSE) | 50 mg |
| Mature stage seed coat (MSSC) | 2-3 mg |

**Figure 1:**
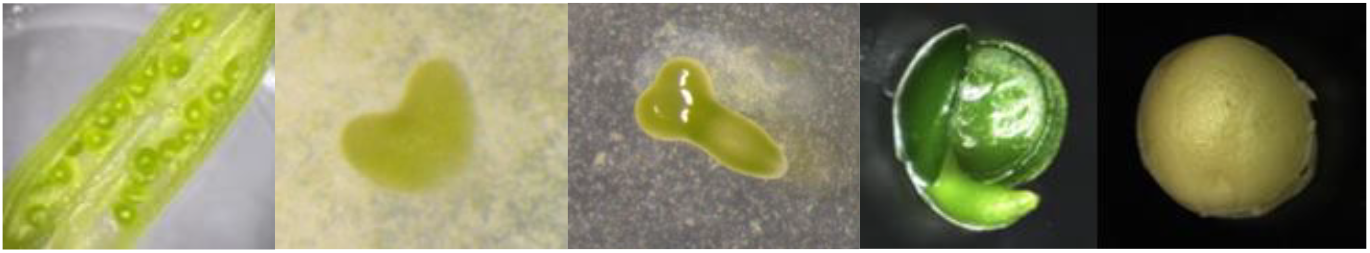
Developmental stages in *Brassica napus* and *B. oleracea*. From left to right: gynoecia wall and ovules, heart stage embryo, torpedo stage embryo, green stage embryo mature stage embryo.

### RNA extraction methods

The RNA from gynoecia walls, ovules, GSE, MSE as well as HSEnd and TSEnd were extracted following the standard E.Z.N.A Plant RNA protocol, including the DNase I (Omega Bio-tek Inc., Norcross, Georgia, USA) digestion protocol on column. Nevertheless, some modifications in the amount of RB buffer and the final elution volume were applied depending on the tissue type. For gynoecia walls and GSE/ MSE, a ratio of 1:12.5 and 1:5 of RB buffer was used, finally eluting the RNA in 20 µl and 30 µl of DEPC water, respectively. The protocol for HSEnd or TSEnd and ovules was slightly modified. For HSEnd and TSEnd, 250 µl of RB with β-mercaptoethanol (β-ME) was added, the tube was vortexed vigorously for 2 minutes allowing the sample to thaw in the RB buffer. For ovules, 150 µl of RB buffer with β-ME was added to the sample to obtain a final volume of 250 µl, before vortexing as before for 2 minutes. The standard E.Z.N.A. Plant RNA kit protocol was then followed, with the elution of RNA in a final volume of 30 µl of DEPC water.

For HSE and TSE, seed coats at all developmental stages and GSEnd, the modifications applied to the Picopure RNA Isolation Kit (ThermoFisher Scientific, Waltham. USA) protocol reported here allowed the extraction of high-quality RNA, with the added benefit of starting with small amounts of seed coat and green endosperm tissue. For seed coats and GSEnd, between 2 and 3 mg of ground tissue was used for RNA extraction. A ratio of 1:2 and 1:3 (v/v) polyvinylpolypyrrolidone (PVPP, ~110 µm particle size, Sigma-Aldrich, Saint Louis, USA) to ground tissue was used, respectively. A 50 µl aliquot of extraction buffer from the Picopure RNA Isolation kit was added, and the samples were homogenized by vortexing for 2 min. Next the samples were incubated for 30 min at 42°C followed by a centrifugation at 2,000 *g* for 2 min to allow the separation of debris and PVPP from the cell extract. This is a critical step, as any impurity can block the column, interfering with the RNA extraction. Subsequent steps follow the protocol described in the Picopure RNA Isolation kit manual, but for clarity the next steps of the protocol are detailed here: the supernatant was transferred to a new 1.5 ml tube, and 50 µl of 70% ethanol was added mixing well by pipetting up and down. The cell extract and ethanol mixture were then transferred to a pre-conditioned RNA purification column (incubated for 5 min with 250 µl of conditioning buffer at room temperature and centrifuged for 1 min at 16,000 *g*). The samples were centrifuged for 2 min at 100 *g*, immediately followed by a centrifugation at 16,000 *g* for 30 s. An aliquot of 100 µl of wash buffer (W1) was added to the purification column followed by a centrifugation of 1 min at 8,000 *g*. An aliquot of 100 µl of wash buffer (W2) was added into the purification column followed by a centrifugation of 1 min at 8,000 *g*. A final wash with 100 µl of wash buffer (W2) was added into the purification column followed by a centrifugation of 2 min at 16,000 *g*. If there was some residual wash buffer in the column, a re-centrifugation for 1 min at 16,000 *g* was performed. After the three washes, the purification column was transferred to a new 1.5 ml tube. An aliquot of 11 µl of elution buffer was pipetted onto the membrane of the purification column, followed by an incubation of 1 min at room temperature. The purification columns were centrifugated for 1 min at 1,000 *g* followed by a final centrifugation for 1 min at 16,000 *g* to elute the RNA. The RNA from three technical replicates were extracted for each sample, combined and treated for DNAse treatments using the TURBO DNA-*free* kit (ThermoFisher Scientific, Waltham, USA). A similar procedure was followed for HSE and TSE without the addition of PVPP. As these tissues were already ground, between 20-25 µl and 10-15 µl of HSE and TSE with RB buffer were used, respectively.

### Determination of RNA quality

The concentration and quality of the RNA was assessed spectrophotometrically at 230, 260 and 280 nm (NanoDrop 1000, LabTech, Heathfield, UK). RNA quality and integrity were evaluated by electrophoresis on 1% agarose gel and the RNA integrity number (RIN) was analysed using an Agilent 2100 Bioanalyzer (Agilent Technologies, Inc.).

## Results

Different RNA extraction protocols are optimal for different tissue types. By using E.Z.N.A Plant RNA kit and modifying the ground tissue amount and the RB buffer ratios as well as the final elution volume, RNA of high-quality and yield was obtained from gynoecia walls, ovules, GSE or MSE and HSEnd and TSEnd (Table 2). However, different methods were tested to extract RNA from HSE or TSE, seed coats and GSend as there are several difficulties encountered when extracting RNA from these tissues. Embryos have a high transcriptional activity, but their small size presents a challenge during grinding for efficient RNA extraction. On the other hand, GSEnd-formed by a residual single layer of cellularised endosperm-and seed coats are transcriptionally less active and contain a lot of contaminants which interfere with RNA extractions. Standard RNA extractions methods using E.Z.N.A. Plant RNA kit, Hot borate combined with RNeasy Plant Mini Kit (Qiagen, Hilden, Germany; Wu et al; 2002) or TRIzol reagent-used for tissues with high lipid content-failed to extract high-quality RNA when attempting to extract RNA from these tissues (Table 3).

**Table 2:** Range of RNA concentration, A260/280 and A260/230 ratios for *B. napus* and *B. oleracea* tissue types extracted with the E.Z.N.A. Plant RNA kit, n=15.

| Tissue | RNA concentration (ng/μl) | A260/280 | A260/230 |
| --- | --- | --- | --- |
| Gynoecia wall | 380.2-827.5 | 2.17-2.21 | 1.88-2.21 |
| Ovules | 158.8-382.7 | 2.10-2.19 | 1.80-2.20 |
| Green stage embryo (GSE) | 749.8-1702.2 | 2.19-2.22 | 1.70-2.24- |
| Mature stage embryo (MSE) | 127.3-639.9 | 2.15-2.21 | 1.94-2.26 |
| Heart stage endosperm (HSEnd) | 108.8-637.9 | 2.17-2.20 | 1.96-2.20 |
| Torpedo stage endosperm (TSEnd) | 102.0-243.4 | 2.18-2.21 | 1.8-2.16 |

**Table 3:** RNA concentration and purity of the RNA obtained from different RNA extraction methods for different *B. napus* and *B. oleracea* tissue types.

| RNA extraction method | Tissue | RNA concentration (ng/μl) | A260/280 | A260/230 |
| --- | --- | --- | --- | --- |
| E.Z.N.A Plant RNA kit | Green stage seed coat (GSSC) | 32.66 | 1.67 | 0.30 |
| TRIzol Reagent | Green stage seed coat (GSSC) | 1.57 | 0.73 | 0.43 |
| Picopure RNA Isolation kit | Green stage seed coat (GSSC) | 113.55 | 2.03 | 1.42 |
| E.Z.N.A Plant RNA kit | Green stage endosperm (GSEnd) | 7.79 | 1.39 | 0.14 |
| Picopure RNA Isolation kit | Green stage endosperm (GSEnd) | 54.64 | 1.80 | 0.58 |
| Picopure RNA Isolation kit | Green stage endosperm (GSEnd)+ PVPP | 147.57 | 2.11 | 1.74 |
| Hot Borate | Mature stage seed coat (MSSC) | 24.15 | 1.62 | 0.47 |
| Picopure RNA Isolation kit | Mature stage seed coat (MSSC) | 53.86 | 2.11 | 0.53 |
| Picopure RNA Isolation kit | Mature stage seed coat (MSSC) + PVPP | 58.27 | 2.02 | 1.59 |
| E.Z.N.A Plant RNA kit | Heart stage embryo (HSE) | 16.16 | 2.30 | 0.10 |
| Picopure RNA Isolation kit | Heart stage embryo (HSE) | 104.10 | 2.07 | 1.97 |
| E.Z.N.A Plant RNA kit | Torpedo stage embryo (TSE) | 91.2 | 2.15 | 2.14 |
| Picopure RNA Isolation kit | Torpedo stage embryo | 218.14 | 2.13 | 2.09 |

Different methods were tested for different tissues. The RNA extraction for green stage seed coat (GSSC) and GSEnd using E.Z.N.A. Plant RNA kit for 50 mg of ground tissue resulted in low A260/280 and A260/230 ratios for both tissues, indicating the presence of contaminants in the samples and low RNA yield for green endosperm. RNA from 100 HSE and 50 TSE was also extracted using E.Z.N.A. Plant RNA kit. For these tissues, the RNA purity was high, but the A260/230 ratio and the RNA concentration were low for HSE but not for TSE. The use of the TRIzol reagent method for GSSC using 50 mg of ground tissue resulted in low yields and low A260/280 and A260/230 ratios. The Hot borate protocol was also tested with 100 mg of mature stage seed coat (MSSC) ground tissue. Although proteinase K was added for the removal of proteins, the RNA obtained presented high levels of polyphenols and polysaccharides contaminants as shown by the low A260/230 ratios. The main constraint of these methods is the high amounts of tissue that are needed for obtaining sufficient high-quality RNA.

The use of the Picopure RNA Isolation kit resulted in better quality RNA for all these tissues. By using aliquots of HSE and TSE, the RNA concentration as well as the A260/280 and A260/230 ratios increased significantly, especially for HSE. For GSEnd and MSSC, using between 2 and 3 mg of ground tissue, higher RNA concentrations and A260/280 and A260/230 ratios were obtained. Due to higher concentration and quality of RNA as well as the small amount of starting material required, the Picopure RNA Isolation kit is a better extraction method for these tissues. This method also worked well for heart and torpedo stages seed coat (HSSC and TSSC, respectively). Inspite of the better quality of the RNA extracted using Picopure RNA Isolation kit, some contaminants were still present in the RNA samples, as indicated by low A260/230 ratios for green endosperm and seed coats. The addition of PVPP reduced the amount of polyphenols significantly in the extracted RNA form these tissues, increasing the A260/230 from 0.2 to 1.2, and for some samples, to 1.8 (Table 3). The resulting yields of total RNA after the DNAse treatment and after using three replicates for the tissues extracted from Picopure RNA Isolation kit were high, as well as the A260/280 and A260/230 ratios, confirming the lack (or low) polysaccharide and protein contamination. Although the A260/230 ratios for GSEnd and seed coat tissues were still on the low side, the integrity of the RNA was confirmed observing clear 28S and 18S ribosomal RNA bands with no RNA degradation and by obtaining RIN values from 7 to 10 (Figure 2, Table 4), confirming the high-quality of the RNA obtained from different *Brassica* seed tissue types.

**Table 4:** Range of RNA Integrity Number (RIN) obtained for different *B. napus* and *B. oleracea* tissue types, n=15.

| Tissue type | RIN |
| --- | --- |
| Ovules | 8-10 |
| Gynoecia wall | 9.9-10 |
| Heart stage embryo (HSE) | 8-9.4 |
| Torpedo stage embryo (TSE) | 7.7-9.3 |
| Green stage embryo (GSE) | 9.9-10 |
| Mature stage embryo (MSE) | 9.7-10 |
| Heart stage endosperm (HSEnd) | 7.3-9.4 |
| Torpedo stage endosperm (TSEnd) | 7-8.4 |
| Green stage endosperm (GSEnd) | 7.8-9.5 |
| Heart stage seed coat (HSSC) | 7-9.4 |
| Torpedo stage seed coat (TSSC) | 7-8.4 |
| Green stage seed coat (GSSC) | 8-9.2 |
| Mature stage seed coat (MSSC) | 7.7-9.3 |

**Figure 2:**
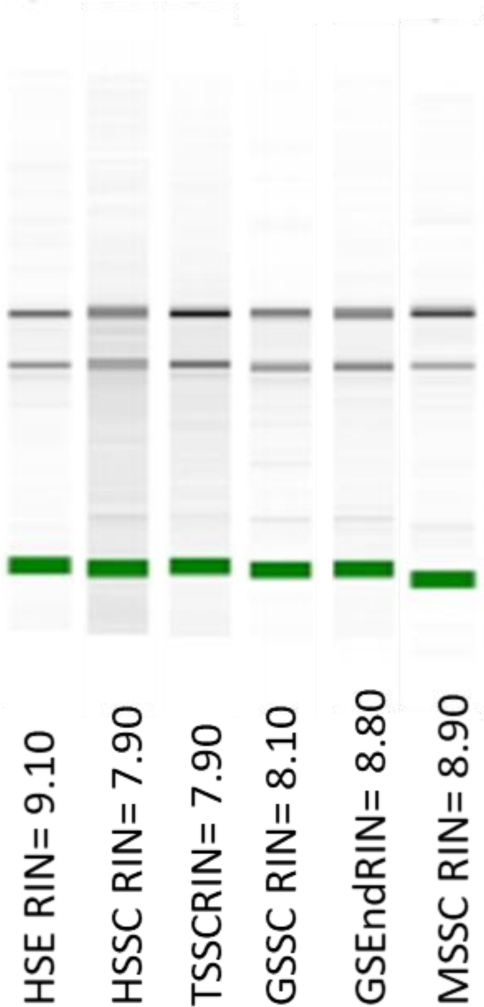
Electropherogram of the RNA isolated from the most challenging seed tissue types. RIN values are shown. HSE: heart stage embryo, HSSC: hear stage seed coat, TSSSC: torpedo stage seed coat, GSC: green stage seed coat, GSEnd: green stage endosperm, MSSC: mature stage seed coat.

## Discussion and conclusions

Obtaining high-quality RNA from different seed tissues can be challenging, especially for tissue types that do not have high transcriptional activity and may contain secondary metabolites as contaminants that interfere with RNA extraction methods. Several protocols were tried to extract good quality RNA from these complex tissues. The TRIzol reagent allows the precipitation of RNA, DNA and proteins from a single sample. However, the resulting yield obtained from 50 mg of GSSC was unexpectedly low. Although the use of the chaotropic agent guanidium isothiocyanate for cell lysis and denaturation of ribonucleases, chloroform to separate RNA from DNA and proteins and isopropanol for RNA precipitation was attempted, the RNA obtained from GSSC using this methodology presented high content of contaminants, as shown by the low A260/280 and A260/230 ratios obtained. For other seed species with high levels of polyphenols, RNA obtained using TRIzol reagent was also degraded and presented low A260/230 ratios (Pereira et al., 2017; Mornkham et al., 2013), suggesting that TRIzol is not a good methodology to extract RNA from seeds or seed coats which contain these secondary metabolites. Although hot borate has been described as a good protocol for extracting RNA from samples rich in polysaccharides, phenolic compounds and oil (Gudenschwager et al., 2012: Wan & Wilkins., 1994), the RNA extracted from MSSC with this method was not pure. MSSC accumulate high amounts of polyphenols in the maturation stage of seed development, specially oxidised procyanidins which confer the dark brown colour of the seed coat characteristic of this stage of development. The low A260/280 and A260/230 ratios obtained from MSSC reflected protein and polyphenols and polysaccharides contamination, respectively. A limitation of the amount of tissue used or the use of proteinase K in this methodology instead of β-Me could be an alternative explanation for the high level of contaminants present in the RNA. Several methods for RNA extraction from whole seeds used PVPP as a removal of polysaccharides, proteins and polyphenol compounds (Kanai et al., 2017; Morkham et al., 2013; Salzman et al., 1999). Here, we confirm that the addition of PVPP to seed coats and GSEnd tissues significantly improved the quality of the extracted RNA by using a ratio of 1:2 or 1:3 of PVPP and ground tissue (v/v). The addition of PVPP in these tissues is important, as in these stages of development *Brassica* spp seeds accumulate polyphenols in the seed coats and polysaccharides in the endosperm. As the 1:2 or 1:3 ratio (v/v) worked for *Brassica* spp seed tissues, the amount of ground tissue needed as well as the PVPP and ground tissue ratio may vary whilst extracting RNA from other species. For other oilseeds, such as *Sesamum indicum*, *Zea mays*, *Helianthus annuus*, *Linum usitatissimum* (Rayani & Nayeri 2015), *Arachis hypogaea* (Dang & Chen, 2012), *Jatropha curcas* (Singh et al., 2009) and *Chamaerops humilis* (Siles et al., 2018), different protocols use a mixture of Tris buffer combined with chloroform:isoamyl alcohol and phenol: chloroform reagents to extract RNA from whole seeds. These methodologies require high amounts of material, from 70 mg to 1 gram. However, the RNA extraction method reported here for HSE, TSE, HSSC, TSSC, GSSC, GSEnd and MSSC using Picopure RNA Isolation kit is a robust protocol for obtaining high-quality RNA for gene expression analyses by only using 2 or 3 mg of tissue and a low number of embryos. Also, the use of β-Me as a reducing agent for denaturing ribonucleases that are released during tissue disruption and homogenization avoided the use of proteinase K, and pure RNA with no signs of degradation was retrieved. Also, the modifications reported here for the E.Z.N.A. Plant RNA kit ended with high yields and pure RNA for more transcriptionally active tissues (gynoecia wall, ovules, HSEnd, TSEnd, GSE and MSE) which present high levels of polysaccharides and lipids. As described by Schroeder et al. (2006); RIN values of 10 indicate completely intact RNA whereas RIN values of 1 indicate totally degraded RNA. The range of high levels of RIN values obtained for different seed tissue types during *Brassica* spp seed development confirmed the high-quality of the extracted RNA using these extraction protocols, being suitable for RNA-Sequencing. In our case, the modifications applied to E.Z.N.A. Plant RNA kit and Picopure RNA Isolation kit extraction methods were successfully used for different *Brassica* spp genotypes including the diploid and polypoid species, obtaining a minimum of 25 million reads and 7.5gb of raw data for all the tissue types.

In conclusion, we report efficient and reliable high-quality RNA extraction methods for different oilseed *Brassica* spp tissue types during seed development and pre-fertilization tissues suitable for RNA-Sequencing and subsequent gene expression analysis.

## Abbreviations

HSE: Heart stage embryo
HSEnd: heart stage endosperm
HSSC: heart stage seed coat
TSE: torpedo stage embryo
TSSC: torpedo stage seed coat
TSEnd: torpedo stage endosperm
GSE: green stage embryo
GSEnd: green stage endosperm
GSSC: green stage seed coat
MSE: mature stage embryo
MSSC: mature stage seed coat

## Acknowledgements

This work was supported by UK Biotechnology and Biological Sciences Research Council grant BB/P003095/1.

---

Supplementary Video 1: Video demonstrating the extraction of liquid HSEnd. The extracted endosperm is shown placed on the glass slide solely for demonstration purpose. This drop would be transferred directly to the microfuge tube upon confirmation of embryo stage.

